# The danger hypothesis of virulence evolution

**DOI:** 10.64898/2026.05.20.726587

**Authors:** Mathias Franz, Roland R. Regoes, Jens Rolff

## Abstract

Multicellular organisms regularly encounter microbes, which are, however, only rarely pathogenic. Our understanding of this phenomenon is currently restricted due to lacking theory on evolutionary transitions between non-pathogenic and pathogenic microbial lifestyles. Here we addressed this gap by investigating a mathematical model of host-microbe interactions that is based on the danger theory of immunology, which states that danger signals related to host tissue damage play a key role in activating immune responses. We formally implemented this idea by assuming that immune activation increases with costs that microbes cause to their host, and we compared this to scenarios in which immune activation depends only on the presence or load of infecting microbes. Our model analysis revealed that cost-based – but not presence or load-based – immune activation favours the evolution of avirulence and associated non-pathogenic microbial lifestyles. Based on our results, we propose the ‘danger hypothesis of virulence evolution’ which states that evolution towards avirulence and intermediate virulence are both possible – depending on whether hosts can accurately assess costs generated by microbes. The idea that basic host immune responses can select for avirulence offers a new explanation for why most microbes are not pathogenic to a given host.

## Introduction

Multicellular organisms regularly encounter microbes, including bacteria, archaea and unicellular eukaryotes (McFall-Ngai et al., 2013). While it has been argued that most of those microbe species are pathogenic to at least one host (Schmid-Hempel, 2021), to a given host only a small proportion of these microbes is pathogenic (Casadevall, 2025; Drake, 2025). This is surprising given that multicellular organisms are potentially very valuable resources for microbes. Current explanations for the small proportion of pathogens stemming from a large pool of rather benign microbial life focus on constraints that make the transition towards pathogenic lifestyle challenging. For example, it has been argued that microbes rarely become pathogens because this requires that microbes acquire unlikely phenotypes that combine many traits, including the virulence factors and physiological characteristics that allow them to infect a host, and thereafter to survive and replicate inside the host (Casadevall, 2025). Here, we offer an alternative explanation based on a mathematical model of microbe virulence evolution. Virulence is here generally defined as reduction in host fitness that is caused by microbes or other parasites (Read, 1994). Specifically, we propose that independently of how frequently pathogenic phenotypes arise, basic host immune responses can select against the evolution of pathogenic lifestyles.

Historically the avirulence hypothesis captured a popular view that pathogenic microbes will eventually evolve towards avirulence, because by harming their hosts, microbes would also reduce their own fitness (Alizon et al., 2009; Méthot, 2012; Smith, 1904). This hypothesis seems to directly explain why only few microbes are pathogenic. However, prevailing theory on virulence evolution – also known as the trade-off hypothesis – rejects the avirulence hypothesis and instead predicts an evolutionary trajectory towards intermediate virulence due to the existence of fitness trade-offs (Acevedo et al., 2019; Alizon et al., 2009; Alizon & Michalakis, 2015; Anderson & May, 1982; Cressler et al., 2016; Ewald, 1983; Frank, 1996; Franz et al., 2025; Leggett et al., 2013). Accordingly, due to the focus on pathogenic microbes, evolutionary transitions between non-pathogenic/avirulent and pathogenic/virulent microbial lifestyles is beyond the scope of current theory on virulence evolution. We addressed this gap by investigating a mathematical model of host-microbe interactions that is based on the danger theory of immunology.

A central concept in immunology is the idea that immune systems of multicellular organisms distinguish between self and non-self, where the detection of non-self indicates the presence of infecting microbes and thus triggers related immune responses (Janeway, 1992; Medzhitov & Janeway Jr, 2002). Based on the immunological self/non-self-theory, previous theoretical work on pathogen virulence evolution, and more generally host-microbe interactions, assumed that immune responses react to the presence or load of infecting microbes (e.g. Alizon & van Baalen, 2005, 2008a; 3 André et al., 2003; Antia et al., 1994; Duneau et al., 2025; Ellner et al., 2021; Gilchrist & Sasaki, 2002; Metcalf & Koskella, 2019).

The self/non-self-theory has been challenged by the danger theory of immunology, which was first proposed by Polly Matzinger (Matzinger, 1994) and subsequently developed further (Gust et al., 2017; Kroemer et al., 2024; Matzinger, 2007). The core argument of this theory states that danger signals, e.g. related to host tissue damage, play a key role in activating immune responses – and related work in immunology focussed on identifying the underlying mechanisms (Kroemer et al., 2024).

Here, we took a more evolutionary perspective on the danger theory of immunology, which we formalized in a mathematical model (Fig. 2). We assumed that danger signals allow hosts to gather information on the costs that infecting microbes cause to the host. Accordingly, rather than implementing specific immunological mechanisms, we assumed that immune reactions respond to microbe-inflicted costs (Fig. 2a). To compare the effects of such danger detection to self/non-self detection, additional scenarios were implemented, in which the immune system only responds to the presence or load of microbes (Fig. 2a). Based on the findings of our model analysis, we developed the ‘danger hypothesis of virulence evolution’, which is based on the danger theory of immunology, and which combines features of the trade-off hypothesis of virulence evolution, and partly also the avirulence hypothesis (Fig. 1).

**Fig. 1:**
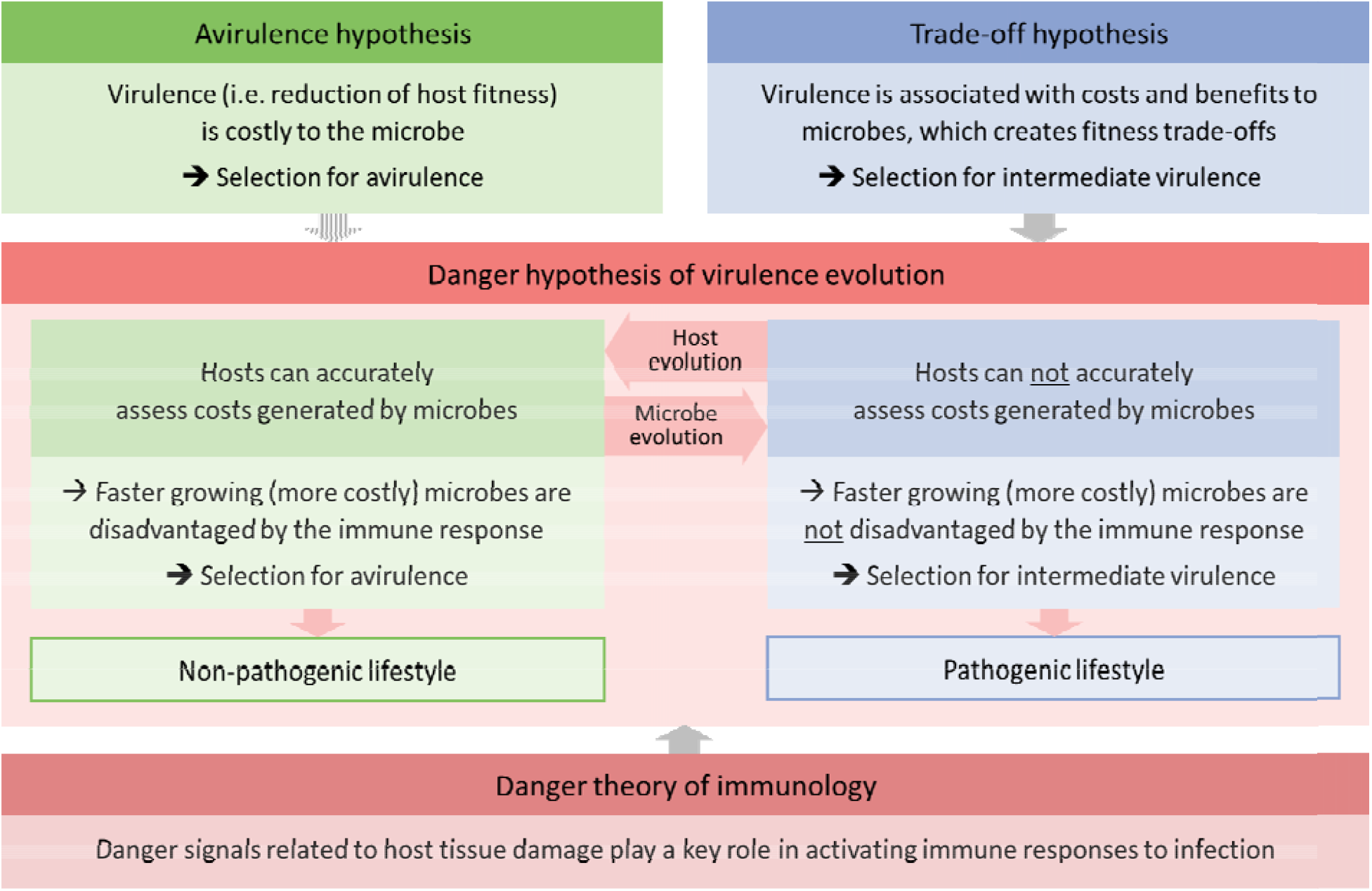
Overview of the proposed ‘danger hypothesis of virulence evolution’ in relation to other theories in evolutionary biology and immunology. Virulence can be generally defined as reduction in host fitness that is caused by microbes or other parasites (Read, 1994). Historically, the avirulence hypothesis had been the main framework for explaining microbe virulence evolution (Alizon et al., 2009; Méthot, 2012; Smith, 1904). However, in the past decades this hypothesis has been replaced by the trade-off hypothesis, which predicts that instead of evolving towards avirulence, pathogenic microbes will evolve towards intermediate virulence (Alizon et al., 2009; Franz et al., 2025). Here we propose the danger hypothesis of virulence evolution, which is based on the danger theory of immunology (Kroemer et al., 2024; Matzinger, 1994), and which combines features of the trade-off hypothesis, and partly also the avirulence hypothesis. In our hypothesis the trajectory of virulence evolution and the resulting microbial lifestyle depends on whether host immune systems can accurately assess costs generated by microbes. This ability of the host immune system is itself subject to host-microbe coevolution, where microbes can evolve means to interfere with the host immune response, and hosts can evolve counter measures. Taken together, our hypothesis provides a new perspective on the evolution of virulence – which includes transitions between non-pathogenic and pathogenic lifestyles for a given host.

**Fig. 2:**
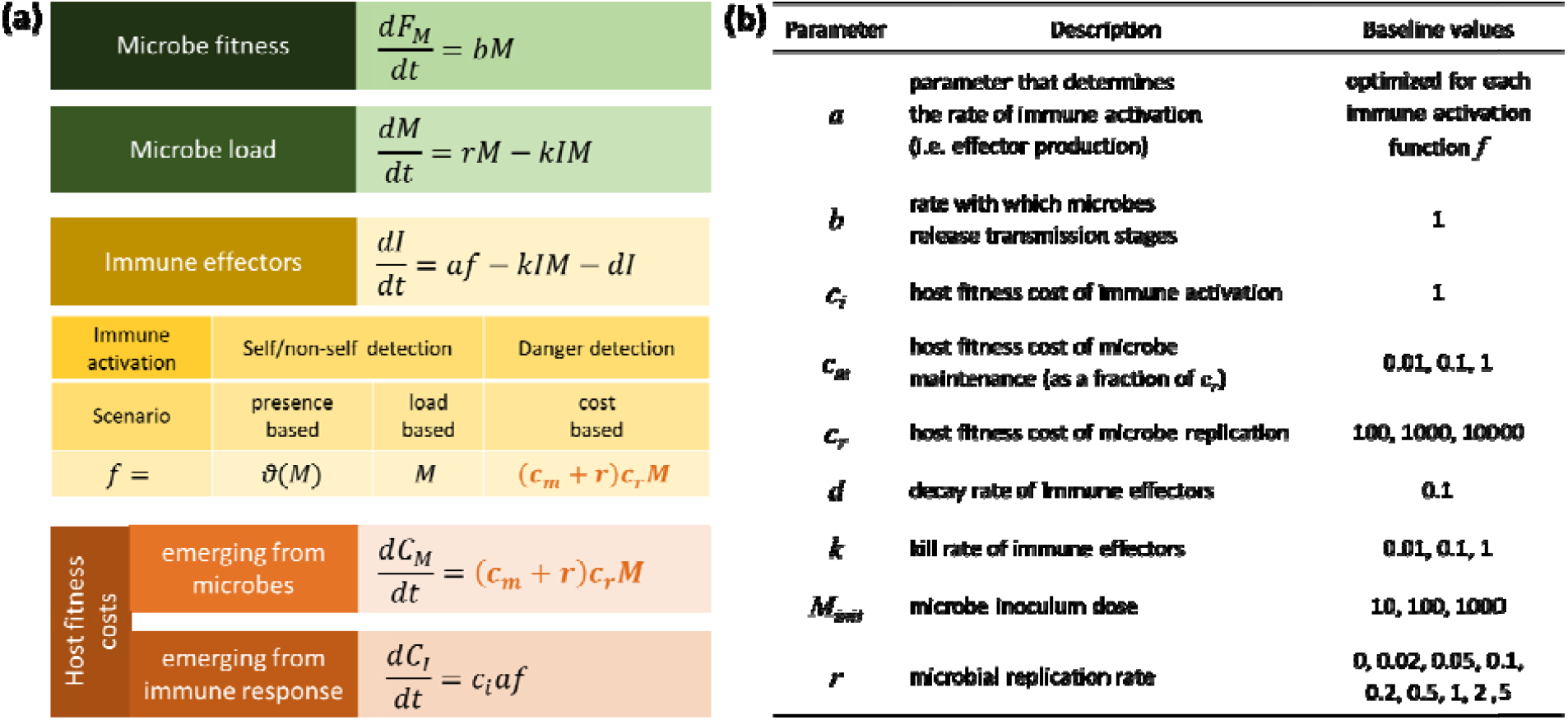
Overview of the mathematical infection model. **(a)** Differential equations describing within-host infection dynamics. Microbe fitness *F*_*M*_ accumulates with the rate *b* by which transmission stages are released from the microbe population *M*. Microbes *M* are growing with rate *r* in absence of an immune response and are killed by immune effectors *I* with rate *k*. In different scenarios, immune effectors are generated based on different immune activation functions *f* and an activation parameter *a*, which is separately optimized for each scenario. While presence-based and load-based scenarios are consistent with self/non-self detection, the cost-based scenario captures danger detection. Degradation of immune effectors occurs during microbe killing and additionally with a constant rate *d*. Abstract host fitness costs accumulate due to costs emerging from microbes (*C*_*M*_) and the host immune response (*C*_*I*_). Microbe-inflicted costs emerge from microbe replication scaled by *c*_*r*_, and microbe maintenance scaled by *c*_*m*_*c*_*r*_. Costs of the immune response emerge from immune activation scaled by *c*_*i*_. An extended model also includes costs related to the quantity of immune effectors (Fig. S5). In the model it is not specified how the abstract fitness costs affect host fitness, but since the possibility of host death is excluded in the model, abstract host fitness cost can be conceived as reducing host fecundity. **(b)** Model parameters and values of the baseline scenario. Main parameters that determine microbe pathogenicity are *c*_*m*_, *c*_*r*_ and *r*. In model simulations, the initial value of *M* was set to *M*_*init*_, and all other state variables (*F*_*M*_, *I, C*_*M*_, *C*_*I*_) were set to 0.

## Results

We implemented host evolution by deriving optimized immune responses that minimize host fitness costs of infections, i.e. combined costs inflicted by microbes and the host immune response. More specifically, we assumed that evolution acts on the strength of the immune activation (parameter *a* in Fig. 2) across infections with different microbes – on a spectrum from minimal to high pathogenicity (see Fig. 2b for assumed parameter ranges). We considered that microbes on the whole spectrum of pathogenicity can infect hosts, e.g. through minor wounds. Accordingly, our model focusses on innate immune responses, which are usually the first line of defence and can be activated very quickly (Schmid-Hempel, 2008). In addition, we exclude the possibility that hosts die during an infection as often, but not always, assumed in models of pathogen virulence evolution (Alizon et al., 2009; Alizon & Michalakis, 2015; Cressler et al., 2016; Franz et al., 2025; Kennedy, 2023). This assumption fits to our focus on the transition between non-pathogenic and pathogenic microbe lifestyles, and facilities the analysis of our model, without, however, restricting our main findings and associated conclusions.

In our analysis, each immune activation scenario was optimized separately, which allowed us to identify the most adaptive scenario in our model. Additionally, the investigation of microbe fitness gradients enabled us to derive implications of detecting self/non-self vs. danger for the evolution of microbe virulence.

### Cost-based immune activation is most adaptive for the host

Our results indicate that the cost-based scenario is the most adaptative due to the lowest average fitness costs (Fig. 3a). A key reason for this outcome is that the cost-based scenario, in contrast to the other scenarios, allows the host to closely match costs directly inflicted by microbes and costs emerging from the immune activation (Fig. 3b).

**Fig. 3:**
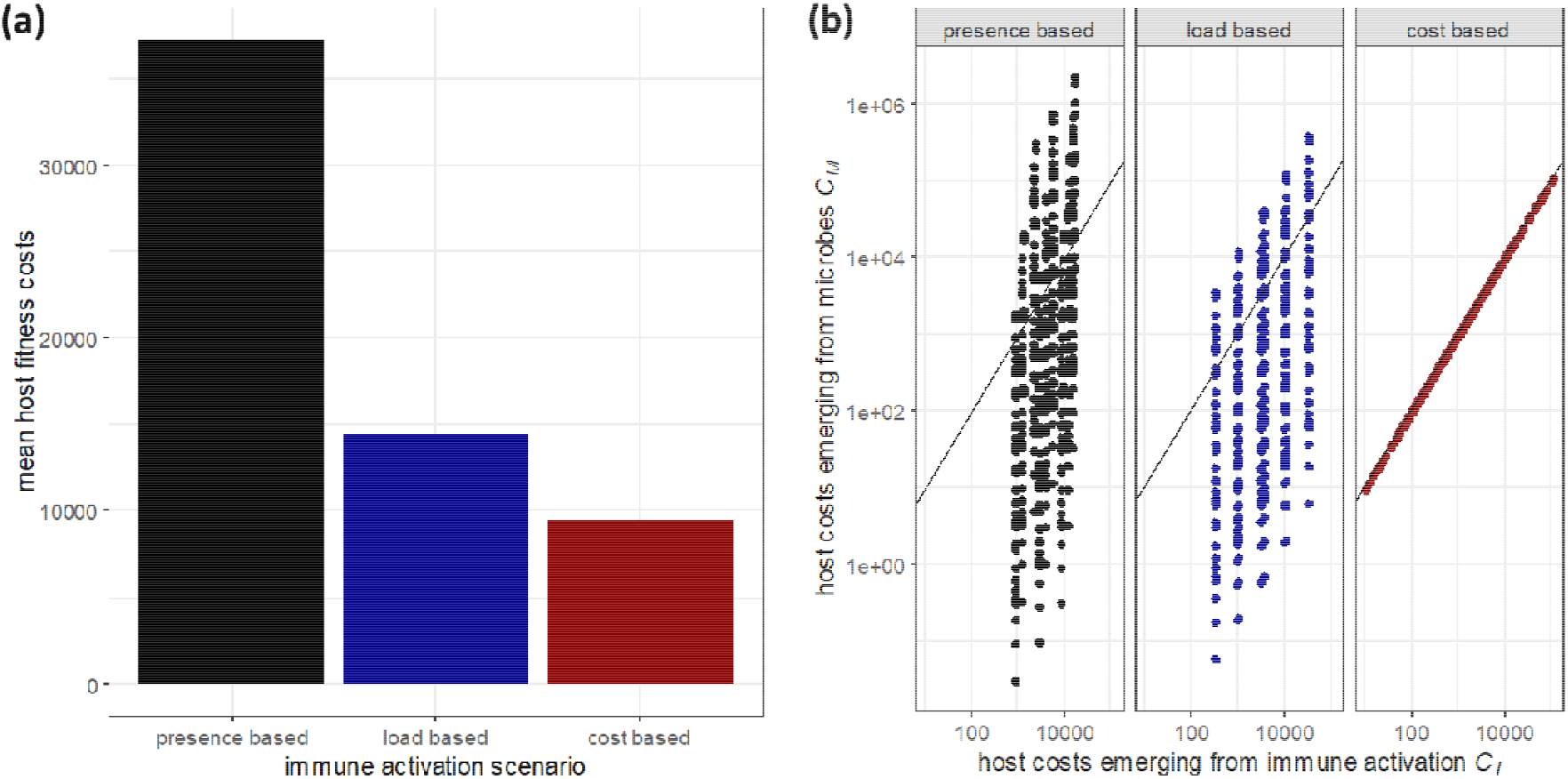
Host fitness costs that emerge during infections with optimized immune activation across a set of microbes with a range of different parameters (see Fig. 2b and Methods section for more details). In the model, host fitness costs are abstract, but can be conceived as reducing host fecundity. **(a)** Mean fitness costs for each immune activation scenario across all simulated parameter combinations. See Fig. S1 for a plot of host fitness costs separately for each parameter combination. **(b)** Costs emerging from the host immune activation vs. costs directly inflicted by the microbe. Each dot represents a unique combination of simulated parameters. The black diagonal line illustrates where both types of costs are equal.

In the presence-based scenario the immune reaction is the least flexible. Accordingly, across infections with different microbes, there is comparably little variation in immune activation costs but large variation in microbe costs (Fig. 3b). In contrast, in the other scenarios, which both rely on additional information about the infecting microbe, there is a larger variation in immune activation costs and smaller variation in microbe costs. Importantly, this reduced variation allows the host to avoid very high microbe costs. This avoidance of very high microbe costs is most pronounced in the cost-based scenario, where microbe and immune activation costs are always closely matched (Fig. 3b). Very similar patterns were obtained 1) in additional analyses in which we varied host specific parameters of immune effector decay rate *d* and costs of immune activation *c*_*i*_ (Fig. S2,S3), and 2) in an extended model in which we included additional costs related to the quantity of immune effectors (Fig. S4).

### Cost based-immune activation selects for avirulent microbes

To investigate the effects of different immune activation scenarios on microbe virulence evolution, we assumed that the fitness of each microbe is proportional to the microbe load during an infection (Fig. 2a, see Methods for more details). Our results for the presence- and load-based immune activation scenarios are fully consistent with current theory on microbe virulence evolution. As expected, increased growth rates resulted in increased microbe fitness (Fig. 4a). The related fitness benefits are rather limited, which is consistent with previous theoretical analyses (Alizon & van Baalen, 2005). In our model, fitness benefits of faster microbial growth arise due to increased mean microbial load during infection (Fig. 4b) and a longer infection duration (Fig. 4c). In addition, we recover the very intuitive effect that increased microbe fitness is associated with increased host fitness costs (Fig. 4d). Taken together, the results for the presence- and load-based scenarios suggest that – in absence of fitness costs, such as increased host death rate, that generate fitness trade-offs – microbes are expected to evolve maximum growth rates that also maximize virulence, i.e. fitness costs for the host.

**Fig. 4:**
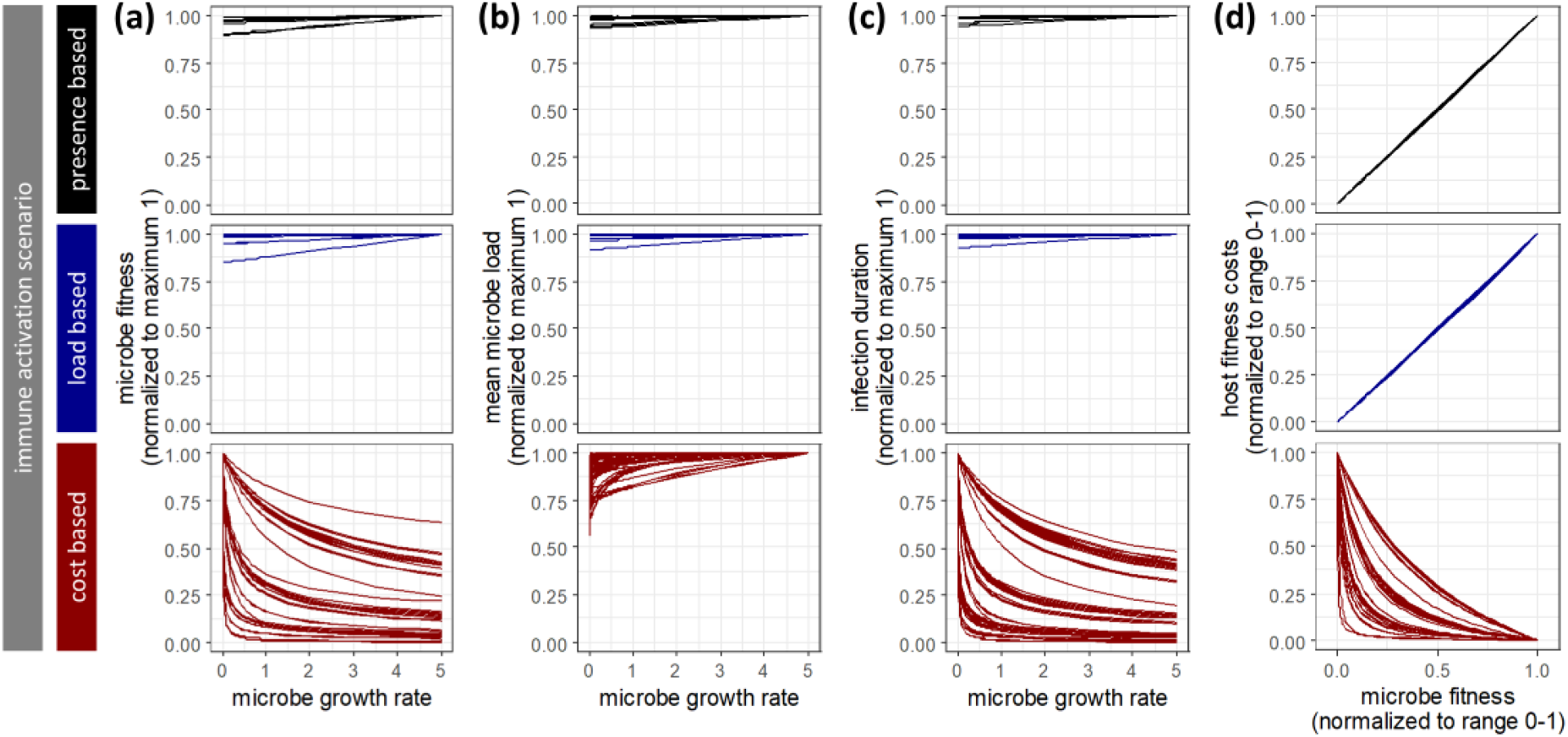
Effects of microbe growth rate variation on microbe and host fitness in different immune activation scenarios. Each line represents a gradient of microbe growth rate *r* for a specific combination of all other varied parameters (Fig. 2b). Accordingly, each line can be interpreted as depicting variants of a microbial strain that differ in growth rates. Results are shown for immune activation parameter *a* that was separately optimized for each scenario. **(a)** Relationship between microbe growth rate *r* and microbe fitness *F*_*M*_, which has been normalized to a maximum of one for each microbe strain. **(b)** Relationship between microbe growth rate *r* and mean microbe load *M* during infection, which has been normalized to a maximum of one for each microbe strain. **(c)** Relationship between microbe growth rate *r* and infection duration, which has been normalized to a maximum of one for each microbe strain. **(d)** Relationship between microbe fitness *F*_*M*_ and host fitness costs (*C*_*M*_+*C*_*I*_), with both values being normalized to a range between zero and one for each microbe strain.

For the cost-based immune activation scenario we obtained very different results. In contrast to general expectation, we found that a higher microbe growth rate led to lower microbial fitness (Fig. 4a). Even though a higher microbial growth rate resulted in increased mean microbial load (Fig. 4b), faster growing microbes were also cleared much faster (Fig. 4c), which explains the growth rate effects on microbial fitness (Fig. 4a). Furthermore, we found the counter-intuitive result that increased microbial fitness was consistently associated with lower host fitness costs (Fig. 4d). Nevertheless, an intuition regarding this relationship can be regained if we accept the possibility that higher microbial fitness arises due to lower growth rates (Fig. 4a), because infections with slower growing microbes trigger a weaker immune response and associated slower clearance of the infection (Fig. 4c). Taken together, the results for the cost-based scenario suggests that – even in absence of additional fitness costs such as increased host death rate – microbes are expected to evolve minimum growth rates that also minimize virulence, i.e. fitness costs for the host.

In addition to effects of microbe growth rates, we also investigated microbe fitness along gradients of microbe growth and maintenance costs (Fig. S6). These analyses revealed that in the cost-based scenario, microbes are expected to evolve minimum growth and maintenance costs that also minimize virulence. To assess the robustness of our findings we conducted additional analyses in which we varied parameters and assumptions related to the host immune response. The results of these analyses (Fig. S7-S9) were consistent with the results reported above.

## Discussion

In this study we used a mathematical model of microbial infections to assess the consequences of immune activation based on self/non-self vs. danger detection (Fig. 2a). We extended previous self/non-self approaches by including a scenario in which immune activation is based on microbe-inflicted costs, which is consistent with the danger theory of immunology (Kroemer et al., 2024; Matzinger, 1994). Our model analysis revealed that this cost-based immune activation scenario outperformed self/non-self scenarios, in which the immune system merely reacts to the presence or number of microbes (Fig 3a). This result emerged because the cost-based immune activation allows the host to closely adjust the strength, and the associated costs, of immune responses to the costs generated by the microbes (Fig. 3b). Accordingly, our model recovers 1) the expectation of immunologists that hosts should escalate immune responses to match the microbial attack (Vance et al., 2009), and 2) the expectation of evolutionary biologists regarding the adaptiveness of danger signals and related information on microbe-inflicted costs (Acuña Hidalgo et al., 2022; Lazzaro & Rolff, 2011). In addition, we found that cost-based immune activation can reverse selection pressures on microbes to favour the evolution of slower microbial growth that is less costly to the host (Fig 4).

Based on our findings, we propose the danger hypothesis of virulence evolution (Fig. 1). This hypothesis translates the mechanistic perspective of the danger theory of immunology into an evolutionary framework, which reconciles the seemingly contradictory ideas of the avirulence and trade-off hypotheses of virulence evolution (Alizon et al., 2009). Evolutionary dynamics expected by these hypotheses, i.e. the evolution of avirulence or intermediate virulence, respectively, can both emerge – depending on whether host immune systems are able to accurately recognize costs generated by the infecting microbe (Fig. 1). Accordingly, through a closer integration of evolutionary and immunological research we provide a new perspective on the evolution of virulence – which includes transitions between non-pathogenic and pathogenic microbial lifestyles.

Although our model supports the idea that microbe evolution towards avirulence is possible under some conditions (Fig. 1, 4a), it is worth noting that the historical avirulence hypothesis (Alizon et al., 2009; Méthot, 2012; Smith, 1904) and our model differ in the underlying cause of avirulence. While in both cases avirulence evolves to avoid a premature end of transmission opportunities, in the historical avirulence hypothesis this is attributed to host death, whereas in our model it is due to microbe clearance by the immune system (Fig. 4c). Importantly, in the cost-based immune activation scenario of our model, faster clearance of more virulent microbes is essentially a byproduct of an immune response that minimizes the costs of an infection to the host (Fig. 3a).

Our hypothesis offers a new answer to the question why most microbes are not pathogenic to a given host (Casadevall, 2025; Drake, 2025). We expect that most microbes encountered by a specific host do not have an existing capacity to interfere with the host immune response. Accordingly, hosts can be expected to mount a cost-based immune response as implemented in our model. In turn, such microbes should experience selection pressures that drive them towards avirulence (Fig. 4a), and thus towards a non-pathogenic lifestyle in relation to the infected host (Fig. 1). However, microbes generally have the capacity to evolve different ways to interfere with the host immune system (Diacovich & Gorvel, 2010; Finlay & McFadden, 2006; Hajishengallis & Lambris, 2011; Van Avondt et al., 2015). If such interference prevents accurate cost estimation, the host immune responses might mainly rely on self/non-self detection, which is captured by the presence and load-based scenarios in our model (Fig. 2). Accordingly, we would expect that such microbes should experience selection pressures that drive them away from avirulence (Fig. 4a), towards a pathogenic lifestyle (Fig. 1). In such cases, additional effects should occur, such as infection-induced host death (Anderson & May, 1982) or sickness behaviour (Ewald, 1983; Kennedy, 2023), which would generate trade-offs that lead to the evolution of intermediate virulence (Alizon et al., 2009; Franz et al., 2025). However, hosts still have the possibility to evolve counter measures to immune interference (e.g. Lopes Fischer et al., 2020), which we expect can re-establish cost-based immune activation and thus revert selection pressures for microbes back towards avirulence (Fig. 1). Taken together, we therefore expect that the reason why microbes rarely become pathogens goes beyond the acquisition of unlikely phenotypes (Casadevall, 2025) and includes evolutionary forces that arise from basic host immune responses, which select against the pathogenic lifestyles (Fig. 1).

Our new hypothesis is expected to be particularly relevant for microbes that form long-term associations with a host, e.g. as in the gut microbiota of many animal species (McFall-Ngai et al., 2013) or mutualistic endosymbionts, which are common in insects (Engel & Moran, 2013; Salem & Kaltenpoth, 2022). The ‘ecosystem on the leash hypothesis’ is well-known for explaining why members of mammalian gut microbiota remain commensals or mutualists – despite a high capacity for evolution and constant opportunities to infect and exploit host tissues (Foster et al., 2017). This hypothesis states that host fitness benefits provided by the gut microbiota lead to the evolution of specific host control mechanisms that disadvantage more pathogenic microbe genotypes. Our hypothesis predicts that such selection pressures should be expected by default even the in absence of specifically evolved control mechanisms — at least if microbes are not interfering with the host immune response (Fig. 1). Thus, our hypothesis provides a solution to the question of how initial host-microbiota association can arise and be stabilized before hosts are able to evolve more elaborate means of microbiota control.

The proposed danger hypothesis of virulence evolution critically rests on the assumption that, if possible, host immune responses to microbe infections should be optimized in a way to minimize the joint fitness costs emerging from microbes and the host immune response. Such an optimization could result purely on the basis of genetic changes in the host. In addition, phenotypic changes during host development might also occur, which could be especially useful for hosts to counter faster microbe evolution. In this context, it has been already documented that early life association of gut microbiota can be important for the maturation of the host immune system and enable appropriate responses to pathogens later in life (Gollwitzer & Marsland, 2015; Weiss et al., 2011). However, the exact mechanism and purpose of this maturation remain to be elucidated. Based on our hypothesis we speculate that an important aspect of such immune maturation could be directed towards improving or fine tuning the accurate recognition of costs generated by infecting microbes.

Current theory related to the trade-off hypothesis of virulence evolution relies heavily on the assumption that microbe virulence evolves due to evolutionary changes in microbe growth rates (Alizon, 2008; Alizon & van Baalen, 2005; André et al., 2003; Antia et al., 1994; Gandon et al., 2002; Ganusov et al., 2002; Gilchrist & Sasaki, 2002). Although, this assumption has been questioned previously (Ebert & Bull, 2003), we are not aware of any studies that directly tested whether changes in microbial growth rates are the main determinant of virulence evolution. For example, in their recent meta-analysis, Acevedo and colleagues (2019) approximated microbe replication with the measure of microbe load during infection. While this approximation is still consistent with less mechanistic trade-off models (Anderson & May, 1982; Day, 2001; Frank, 1996), it remains unclear whether evolutionary changes in virulence are indeed related to microbe growth rates – or whether they are potentially related to variation in immune interference (Diacovich & Gorvel, 2010; Finlay & McFadden, 2006; Hajishengallis & Lambris, 2011; Van Avondt et al., 2015). Although there has been some initial work to assess the effects of immune interference on virulence evolution (Kamiya et al., 2018; Schmid-Hempel, 2008), there seems to be only limited knowledge regarding how variation in immune interference affects microbe fitness gradients and associated evolution of virulence. Pursuing these questions in future theoretical and empirical studies would be very valuable for testing and further developing the proposed danger hypothesis of virulence evolution. While here we focussed on single cell organisms, our arguments likely apply to the virulence evolution of multicellular pathogens.

## Methods

Our model is based on differential equations that describe the population dynamics of a microbe inside a host, the host immune effectors and the fitness consequences for the host and the microbe (Fig. 2). Specifically, we assumed that microbes *M* are growing with rate *r* in absence of an immune response. Immune effectors *I* kill microbes with rate *k*, and are generated based on different immune activation functions *f*. As a baseline scenario we assumed immune activation based on the presence of microbes. In addition, following assumptions of existing virulence evolution models, we considered a load-based scenario in which the number of microbes determines immune activation (e.g. Alizon & van Baalen, 2005, 2008b; Gilchrist & Sasaki, 2002). Finally, to capture dynamics expected according to the danger theory of immunology (Kroemer et al., 2024; Matzinger, 1994), we considered a cost-based scenario in which the immune activation increases with the costs that are generated by microbes at given point during the infection. For each of these scenarios, we assumed that the strength of the immune activation is scaled by an activation parameter *a*, which is separately optimized for each scenario. Degradation of immune effectors occurs during microbe killing and additionally with a constant rate *d*.

We assumed that abstract fitness costs for the host accumulate due to costs emerging from the microbe *C*_*M*_ and also from the host immune response *C*_*I*_. Here, we did not specify how these abstract costs affect host fitness, but since we excluded host death during infection, fitness costs can be assumed to reduce host fecundity. On the microbe side we considered that costs *C*_*M*_ are directly related to microbe replication, which are determined by the parameter *c*_*r*_. In addition, we assumed maintenance costs in absence of replication, which are determined by the parameter *c*_*m*_. More specifically, maintenance costs were assumed to equal a fraction of replication costs, and accordingly maintenance costs were assumed to equal the product of *c*_*m*_ and *c*_*r*_. For each cost type, we assumed that costs can relate to loss of host nutrients as well as any form of damage to the host, which allows the microbe to access nutrients and facilitate microbe replication or maintenance. On the host side, costs *C*_*I*_ were assumed to be proportional to the amount of immune activation, and be determined by parameter *c*_*i*_. In a sensitivity analysis presented in the electronic supplementary materials, we included additional costs based on the quantity of immune effectors that are present at a given point in the infection, which could capture effects of immunopathology (Graham et al., 2005).

Microbe fitness *F*_*M*_ was assumed to be proportional to the microbe load during the course of the infection (Antia et al., 1994). Specifically, we assumed that microbes release transmission stages with rate b. In our model we excluded the possibility that hosts can die during the infection. Accordingly, we excluded the commonly assumed effect that microbe fitness can be reduced by more virulent microbes, which elevate host death rate and thus shorten the duration of the infection (Alizon & van Baalen, 2005, 2008b; Anderson & May, 1982; Day, 2001; Day et al., 2007; Gilchrist & Coombs, 2006; Gilchrist & Sasaki, 2002). We specifically, chose to exclude this possibility, to facilitate the comparison of different immune activation scenarios.

We implemented and numerically analysed our model in the statistical software R (Team, 2022) using the package deSolve (Soetaert et al., 2010). To initialize simulations, we set all state variables to zero except for initial microbe load, which was determined by parameter *M*_*init*_. In case microbe density fell below a value of one, infections were assumed to end (similar to assumptions made by André et al. (2003)). Our main analysis was conducted based on parameter values listed in Fig. 2b. Notably, we assumed that the distribution of growth rates among microbes is skewed towards lower values. We assumed large ranges of kill rates, inoculum doses and all cost parameters. Values *c*_*m*_ were chosen so that maintenance costs were always smaller or equal to replication costs. Costs of immune activation were chosen so that the costs of generating enough immune effectors to kill one microbe (*c*_*i*_/*k*) were always smaller than the cost one microbe generates when it replicates (*c*_*r*_).

For each immune activation scenario, we optimized the immune activation parameter *a* based on simulations for all possible microbe parameter combinations (n=729, with specific parameter values being listed in Fig. 2b). For each parameter combination and given value of *a* we calculated the total host fitness costs (*C*_*M*_+*C*_*I*_) at the end of each infection and used the *optim* function in R to derive the value of *a* that minimizes these costs across infection with all microbes. Subsequently, we simulated infection dynamics for each immune activation scenario with the correspondingly optimized parameter *a* to assess how scenarios affect host fitness costs and pathogen fitness gradients. In addition, we ran a sensitivity analysis to assess the robustness of our results when changing host specific parameters of immune effector decay rate *d* and costs of immune activation *c*_*i*_. Furthermore, we investigated an extended model in which we considered that the host immune response can generate additional costs, which are based on the quantity of immune effectors that are present at a given point in the infection (Fig. S5).

## Acknowledgements

We thank Dino McMahon, Jessica Metcalf and Adam Dobson for helpful feedback on a previous version of the manuscript. This work was supported by the Deutsche Forschungsgemeinschaft DFG: funding to M.F. (grant no. FR 3061/6-1), Research Unit FOR 5026 ‘InsectInfect’ to JR and Mercator fellowship to R.R.R.

