## Supplementary Materials for "The danger hypothesis of virulence evolution"

### Electronic supplementary materials for:

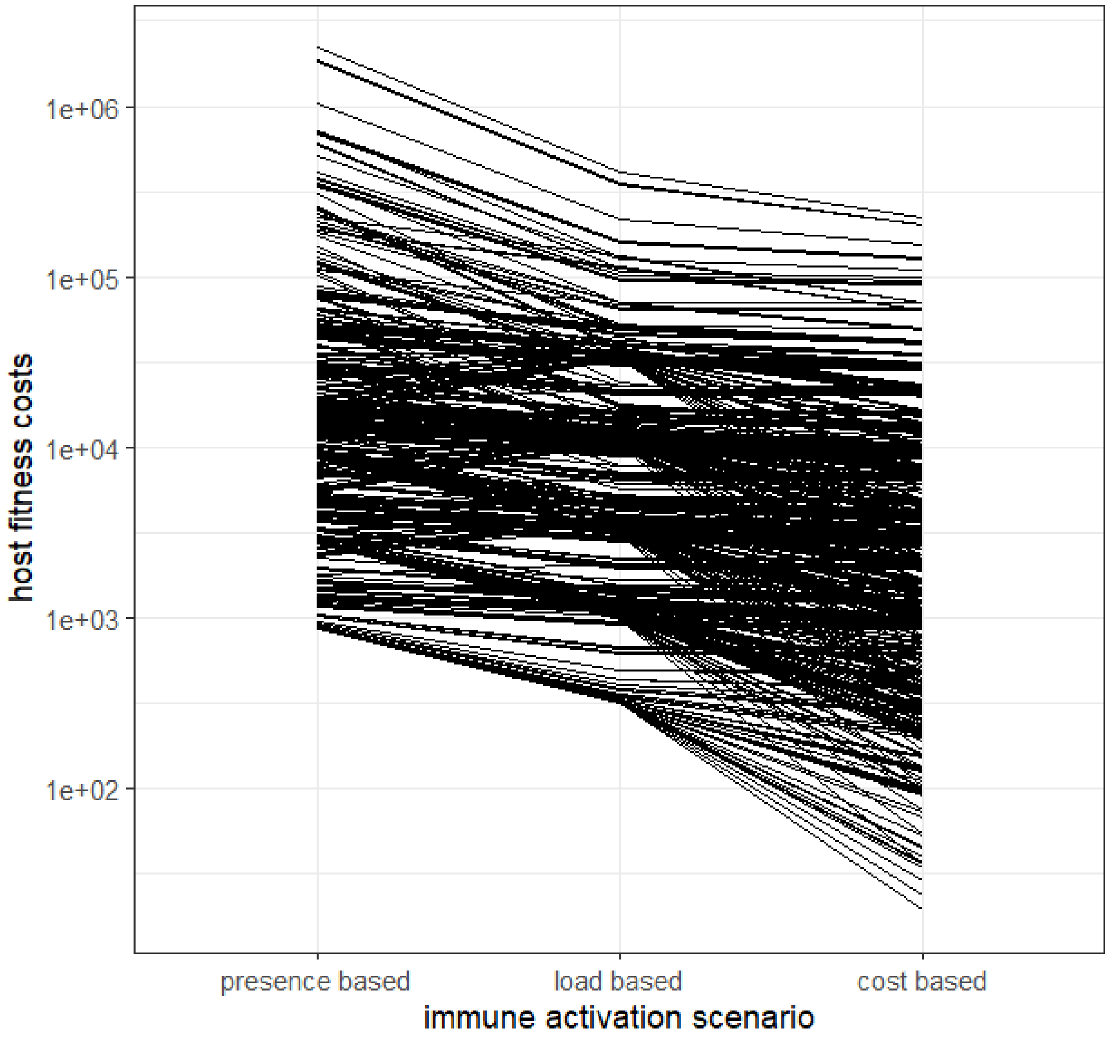


**Fig. S1:** Host fitness costs for each immune activation scenario, separately for each parameter combination.

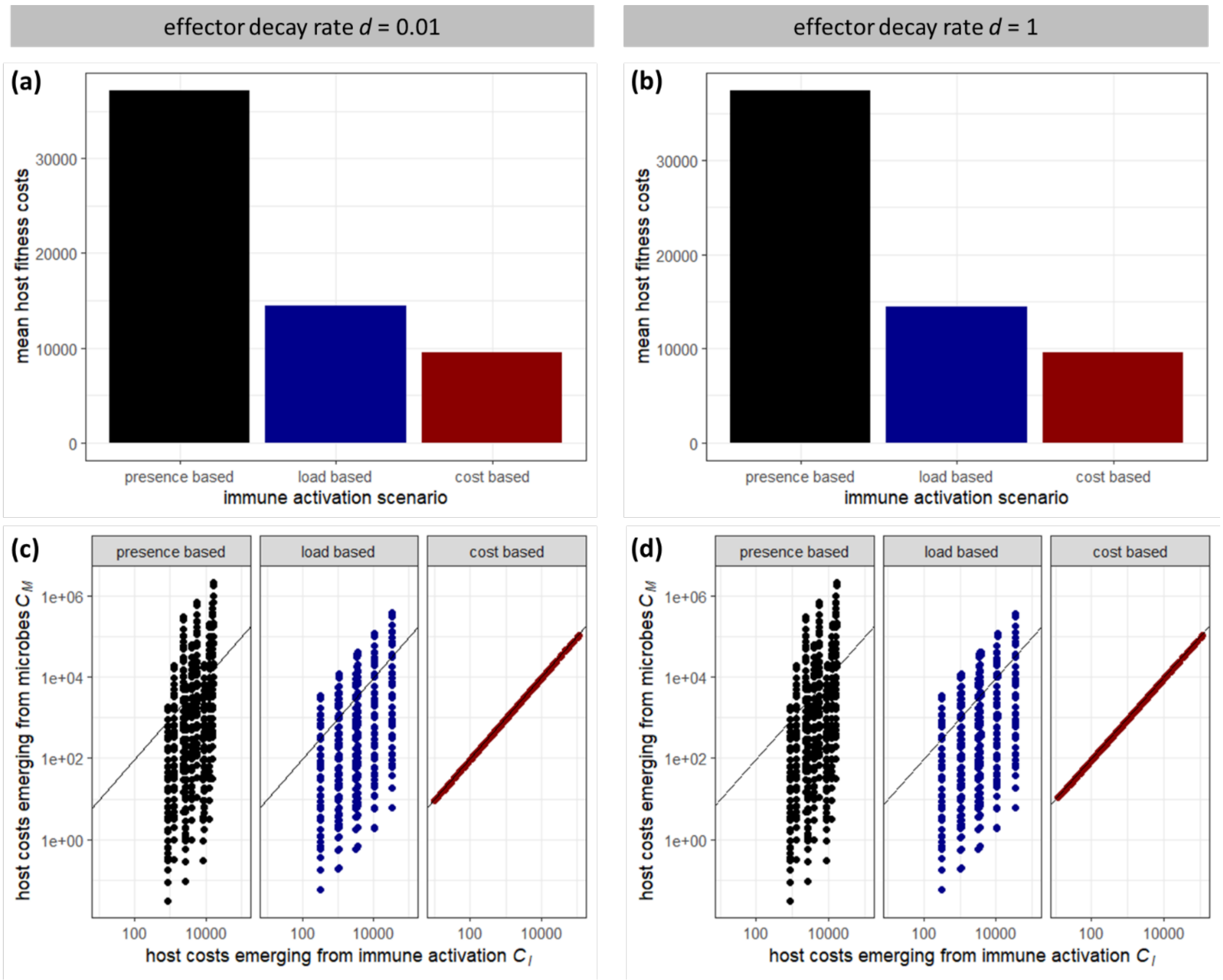


**Fig. S2:** Host fitness costs during infections with optimized immune activation for different effector decay rates *d*. **(a,b)** Mean fitness costs for each immune activation scenario across all simulated parameter combinations. Baseline values (Fig. 1b) were assumed for all parameters except for decay rates *d*. **(c,d)** Costs emerging from the host immune activation vs. costs emerging from the microbe, with each dot representing a unique combination of simulated parameters. The black diagonal line illustrates where both types of costs are equal.

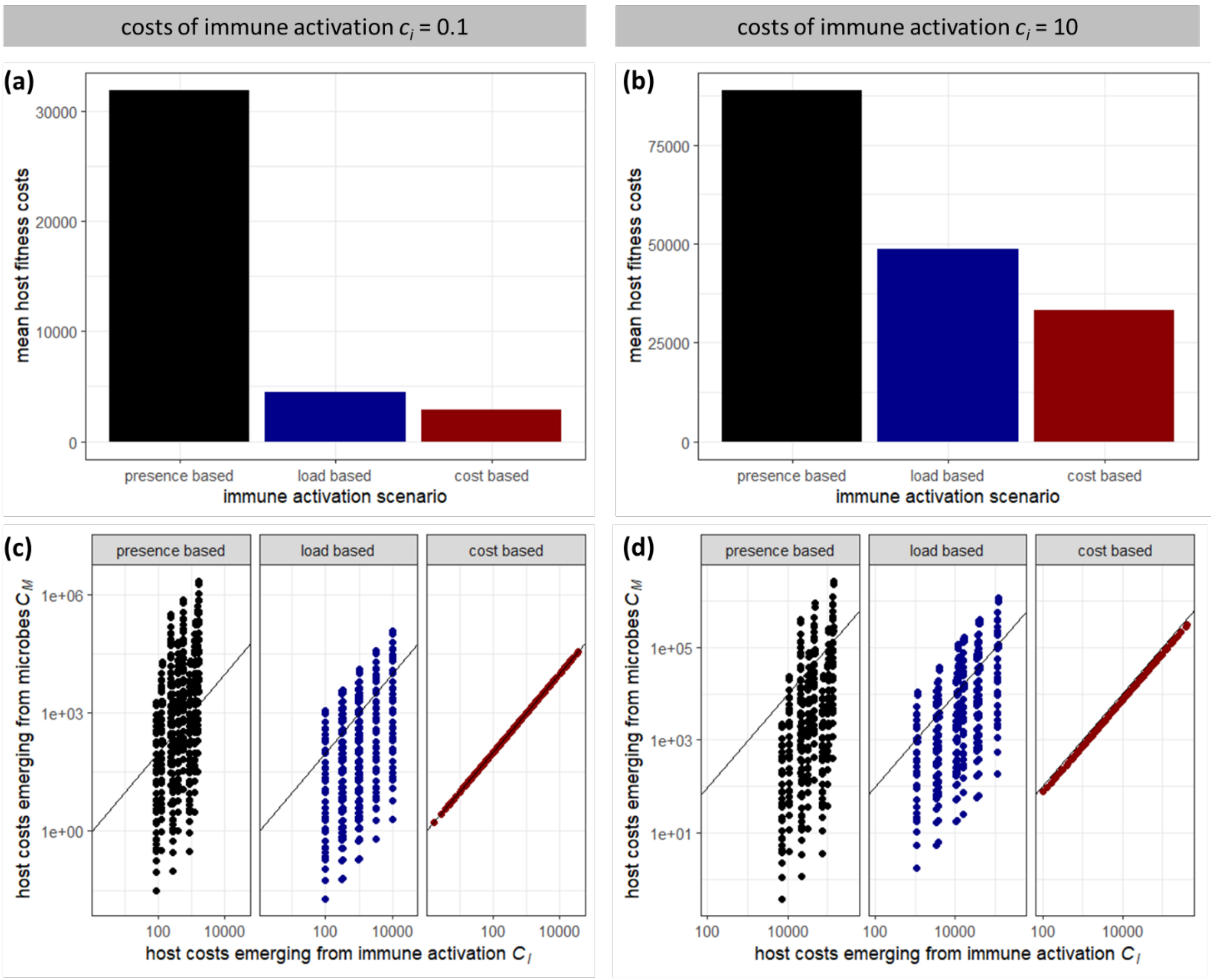


**Fig. S3:** Host fitness costs during infections with optimized immune activation for different costs of immune activation $c_i$. **(a,b)** Mean fitness costs for each immune activation scenario across all simulated parameter combinations. Baseline values (Fig. 1b) were assumed for all parameters except for immune activation $c_i$. In addition, for $c_i$ = 10, values of $c_r$ were restricted to 1000 and 10000 (i.e. the value of 100 was removed) to avoid conditions in which the costs of generating enough immune effectors to kill one microbe ($c_i/k$) are larger than the costs one microbe generates when it replicates ($c_r$). **(c,d)** Costs emerging from the host immune activation vs. costs emerging from the microbe, with each dot representing a unique combination of simulated parameters. The black diagonal line illustrates where both types of costs are equal.

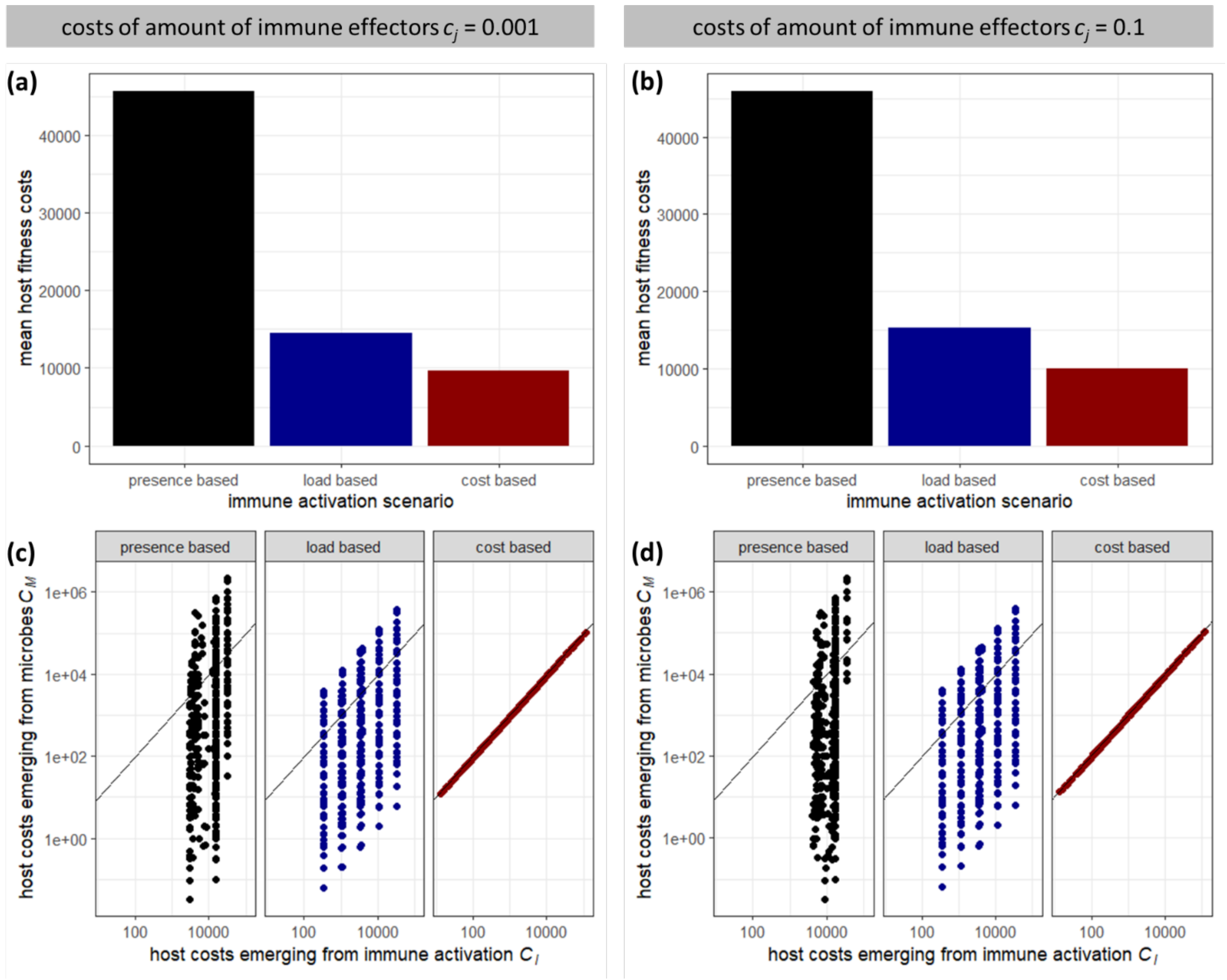


**Fig. S4:** Host fitness costs during infections with optimized immune activation for different costs related to the quantity of immune effectors $c_j$ (see Fig. S5 for an extended model that includes such costs). Baseline values (Fig. 1b) were assumed for all parameters except for $c_j$, and d, which was set to 1 to achieve faster reduction of immune effectors after microbe clearance. All simulations were run for 100 time units. **(a,b)** Mean fitness costs for each immune activation scenario across all simulated parameter combinations. **(c,d)** Costs emerging from the host immune activation vs. costs emerging from the microbe, with each dot representing a unique combination of simulated parameters. The black diagonal line illustrates where both types of costs are equal.

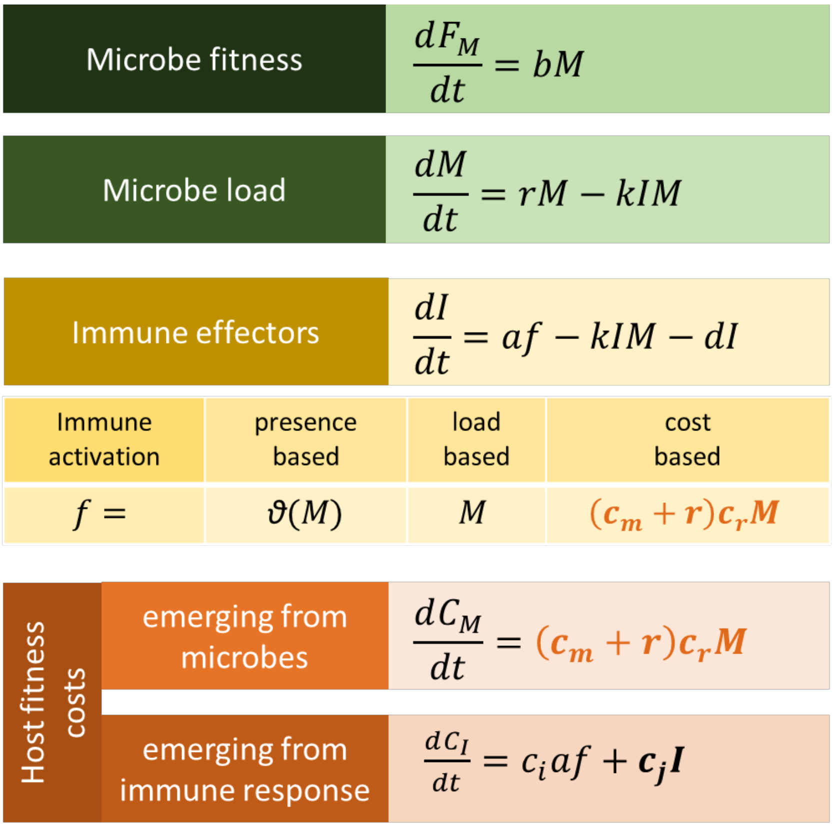


**Fig. S5:** Extended model in which host fitness costs emerging from the immune response include costs $c_j$ related to the quantity of immune effectors that are present at a given point in the infection.

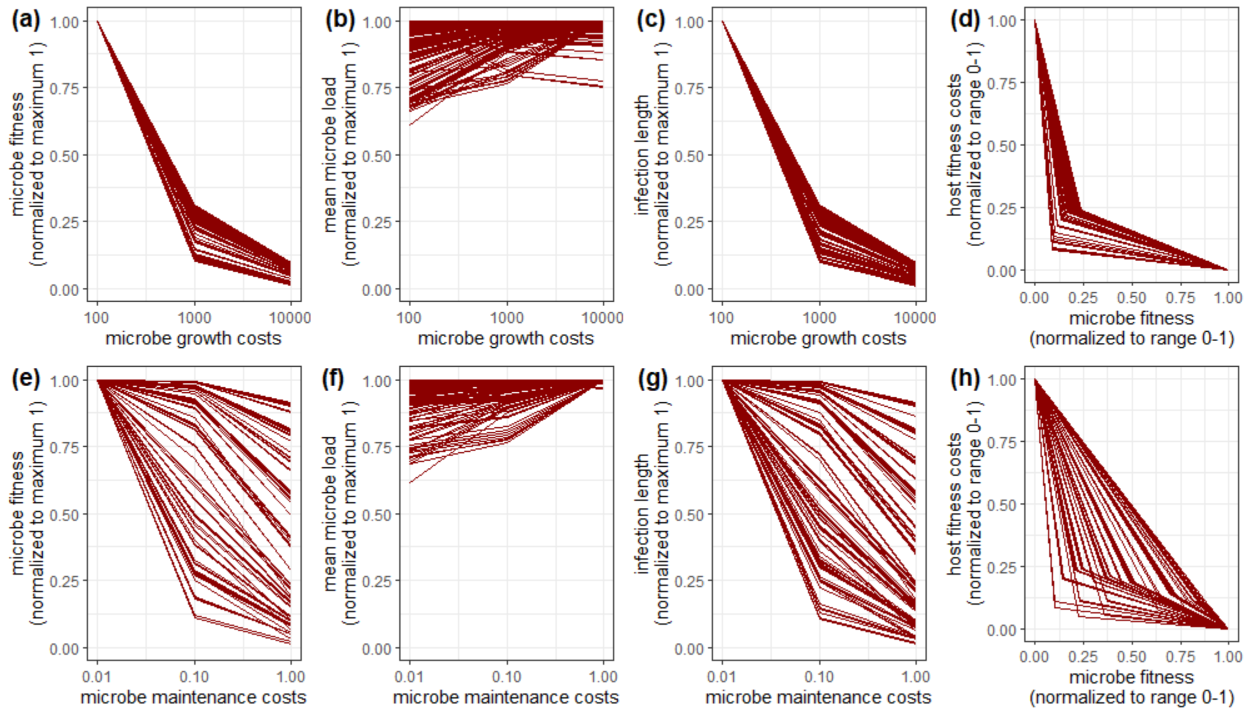


**Fig. S6:** Effects of variation in microbe costs parameters on microbe and host fitness in the cost-based immune activation scenario. Results for other immune activation scenarios are omitted here because variation in microbe cost parameters does not affect microbe fitness in these scenarios. **(a-d)** Effects of microbe growth costs $c_r$ with each line representing a gradient of $c_r$ for a specific combination of all other varied parameters (Fig. 1b). Accordingly, each line can be interpreted as depicting variants of a microbial strain that differ in microbe growth costs. **(e-h)** Effects of microbe maintenance costs $c_m$ with each line representing a gradient of $c_m$ for a specific combination of all other varied parameters (Fig. 1b). Accordingly, each line can be interpreted as depicting variants of a microbial strain that differ in microbe maintenance costs. **(a,e)** Relationship between microbe costs and microbe fitness $F_M$, which has been normalized to a maximum of one for each microbe strain. **(b,f)** Relationship between microbe costs and mean microbe load during infection, which has been normalized to a maximum of one for each microbe strain. **(c,g)** Relationship between microbe costs and infection length, which has been normalized to a maximum of one for each microbe strain. **(d,h)** Relationship between microbe fitness and host fitness costs, with both values being normalized to a range between zero and one for each microbe strain.

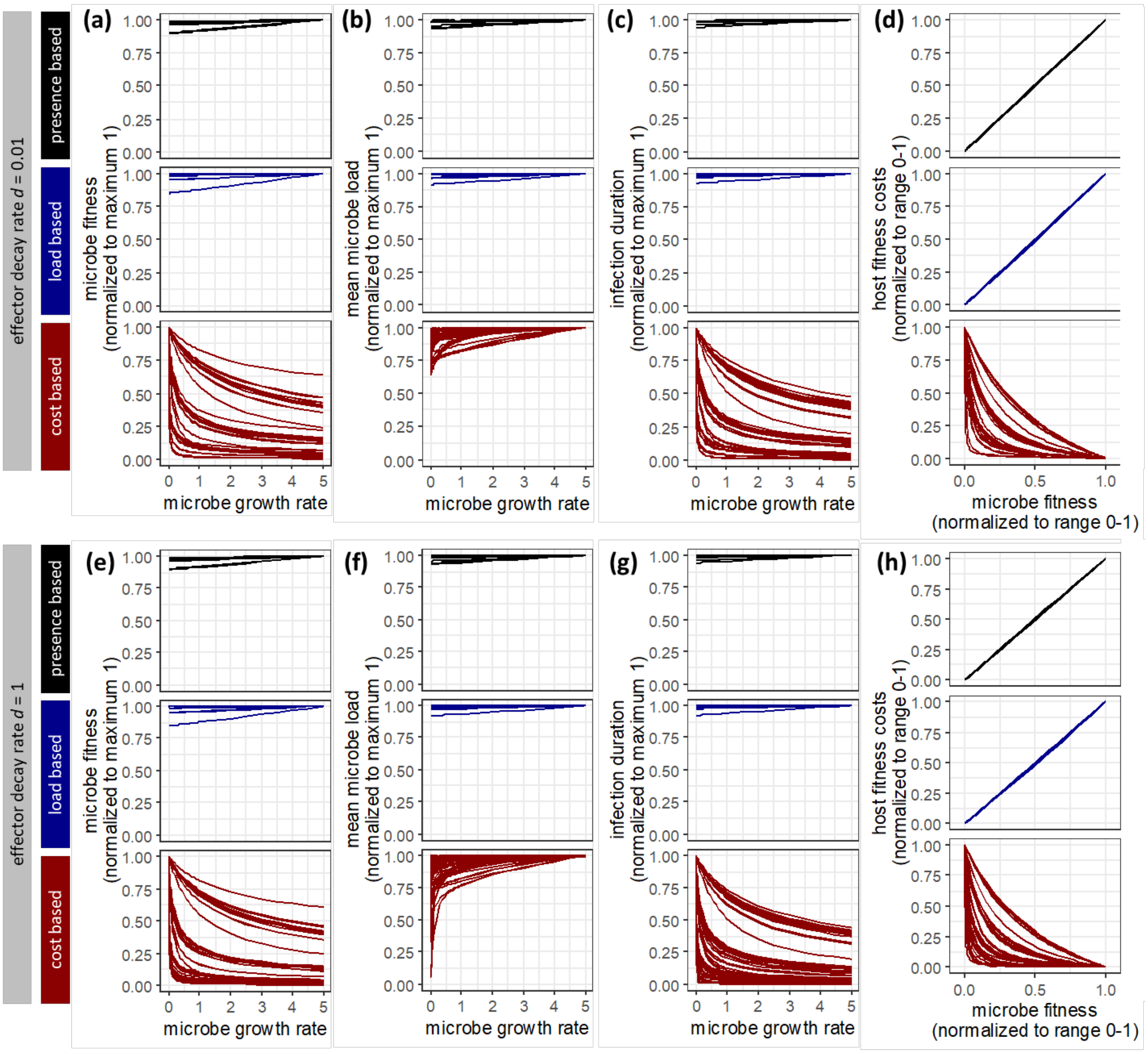


**Fig. S7:** Effects of microbe growth rate variation on microbe and host fitness in different immune activation scenarios and for different effector decay rates *d*. Each line represents a gradient of microbe growth rate *r*. Baseline values (Fig. 1b) were assumed for all parameters except for decay rates *d*. Results are shown for immune activation parameter *a* that were optimized for each scenario. **(a)** Relationship between microbe growth rate *r* and microbe fitness $F_M$, which has been normalized to a maximum of one for each microbe strain. **(b)** Relationship between microbe growth rate *r* and mean microbe load *M* during infection, which has been normalized to a maximum of one for each microbe strain. **(c)** Relationship between microbe growth rate *r* and infection duration, which has been normalized to a maximum of one for each microbe strain. **(d)** Relationship between microbe fitness and host fitness costs, with both values being normalized to a range between zero and one for each microbe strain.

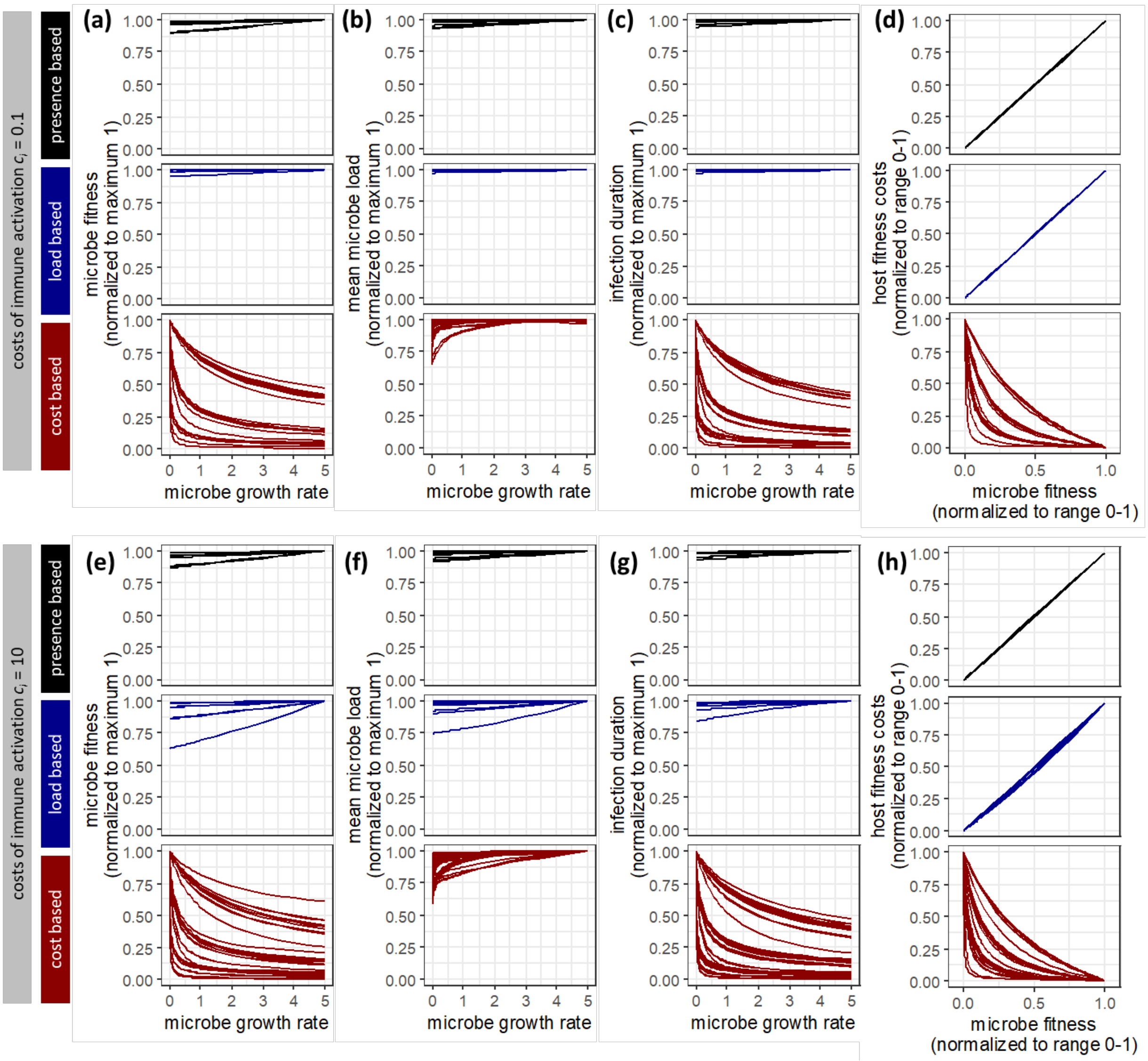


**Fig. S8:** Effects of microbe growth rate variation on microbe and host fitness in different immune activation scenarios and for different costs of immune activation $c_i$. Each line represents a gradient of microbe growth rate $r$. Baseline values (Fig. 1b) were assumed for all parameters except for costs of immune activation $c_i$. In addition, for $c_i$ = 10, values of $c_r$ were restricted to 1000 and 10000 (i.e. the value of 100 was removed) to avoid conditions in which the costs of generating enough immune effectors to kill one microbe ($c_i/k$) are larger than the costs one microbe generates when it replicates ($c_r$). Results are shown for immune activation parameter $a$ that were optimized for each scenario. **(a)** Relationship between microbe growth rate $r$ and microbe fitness $F_M$, which has been normalized to a maximum of one for each microbe strain. **(b)** Relationship between microbe growth rate $r$ and mean microbe load $M$ during infection, which has been normalized to a maximum of one for each microbe strain. **(c)** Relationship between microbe growth rate $r$ and infection duration, which has been normalized to a maximum of one for each microbe strain. **(d)** Relationship between microbe fitness and host fitness costs, with both values being normalized to a range between zero and one for each microbe strain.

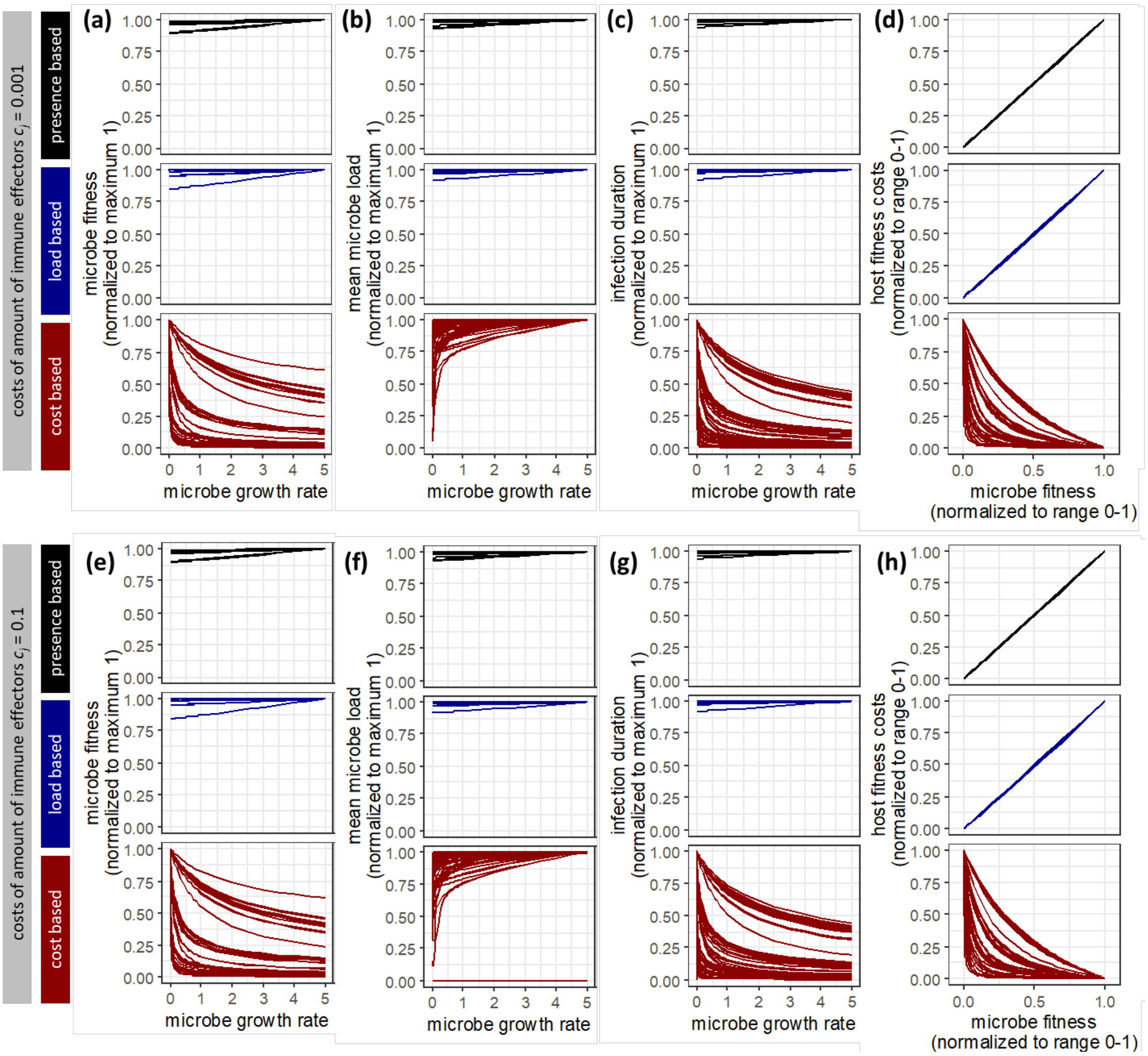


**Fig. S9:** Effects of microbe growth rate variation on microbe and host fitness in different immune activation scenarios and for different costs related to the quantity of immune effectors $c_j$ (see Fig. S5 for an extended model that includes such costs). Each line represents a gradient of microbe growth rate $r$. Baseline values (Fig. 1b) were assumed for all parameters except for $c_j$, and d, which was set to 1 to achieve faster reduction of immune effectors after microbe clearance. All simulations were run for 100 time units. Results are shown for immune activation parameter $a$ that were optimized for each scenario. **(a)** Relationship between microbe growth rate $r$ and microbe fitness $F_M$, which has been normalized to a maximum of one for each microbe strain. **(b)** Relationship between microbe growth rate $r$ and mean microbe load $M$ during infection, which has been normalized to a maximum of one for each microbe strain. **(c)** Relationship between microbe growth rate $r$ and infection duration, which has been normalized to a maximum of one for each microbe strain. **(d)** Relationship between microbe fitness and host fitness costs, with both values being normalized to a range between zero and one for each microbe strain.
